# The Mammalian Cytosolic Thioredoxin Reductase Pathway Acts via a Membrane Protein to Reduce ER-localised Proteins

**DOI:** 10.1101/830026

**Authors:** Xiaofei Cao, Sergio Lilla, Zhenbo Cao, Marie Anne Pringle, Ojore BV Oka, Philip J Robinson, Tomasz Szmaja, Marcel van Lith, Sara Zanivan, Neil J Bulleid

## Abstract

Folding of proteins entering the mammalian secretory pathway requires the insertion of the correct disulfide bonds. Disulfide formation involves both an oxidative pathway for their insertion and a reductive pathway to remove incorrectly formed disulfides. Reduction of these disulfides is critical for correct folding and degradation of misfolded proteins. Previously, we showed that the reductive pathway is driven by NADPH generated in the cytosol. Here, by reconstituting the pathway using purified proteins and ER microsomal membranes, we demonstrate that the thioredoxin reductase system provides the minimal cytosolic components required for reducing proteins within the ER lumen. In particular, saturation of the pathway and its protease sensitivity demonstrates the requirement for a membrane protein to shuttle electrons from the cytosol to the ER lumen. These results provide compelling evidence for the critical role of the cytosol in regulating ER redox homeostasis to ensure correct protein folding and to facilitate the degradation of misfolded ER proteins.

## Introduction

Proteins entering the secretory pathway undergo several modifications that are unique to the ER including glycosylation and disulfide formation (Braakman & Bulleid, 2011). The consequence of these modifications is often increased stability of the protein fold in preparation for secretion into the extracellular milieu. Failure to fulfil the correct modification can lead to protein misfolding and disease (Wang & Kaufman, 2016). Correct protein disulfide formation requires resident ER proteins to catalyse both disulfide formation and reduction (Ellgaard, Sevier et al., 2018). Disulfide formation is catalysed primarily by Ero1 which oxidises members of the protein disulfide isomerase (PDI) family by coupling the reduction of oxygen to the introduction of a disulfide (Sevier, Qu et al., 2007). PDI then exchanges its disulfide with substrate proteins during and following their translocation into the ER lumen (Chen, Helenius et al., 1995). During this process, disulfides may form that are not present in the final native structure (Jansens, van Duijn et al., 2002). Such non-native disulfides need to be removed to allow correct folding or to facilitate protein degradation (Ushioda, Hoseki et al., 2008). Hence, there is a requirement for both an oxidative and reductive pathway in the ER to ensure correct, native disulfides are formed or for proteins to be targeted for degradation (Bulleid & Ellgaard, 2011). In addition to the reduction of structural disulfides, there is also a requirement for a reductive pathway to recycle enzymes such as methionine sulfoxide reductase (Cao, Mitchell et al., 2018), vitamin K epoxide reductase (Rishavy, Usubalieva et al., 2011), peroxiredoxin IV (Tavender, Springate et al., 2010) and formyl glycine generating enzyme (Dierks, Dickmanns et al., 2005) that contain active site thiols which become oxidised during catalysis.

Despite our understanding of the various pathways to introduce disulfides, our knowledge of the reductive pathway is limited. Members of the PDI family such as ERdj5 are likely to catalyse the initial stage in the reduction of non-native disulfides (Jessop, Chakravarthi et al., 2007, Oka, Pringle et al., 2013, Ushioda et al., 2008). ERdj5 has a relatively low reduction potential making it an efficient reductase (Hagiwara, Maegawa et al., 2011). ERdj5 is recruited to substrate proteins via its interaction with BiP (Cunnea, Miranda-Vizuete et al., 2003, Hagiwara et al., 2011, Oliver, van der Wal et al., 1997). Following the reduction of substrates, the PDI reductases will become oxidised so to maintain activity there is a requirement for their enzymatic reduction. Within the cytosol, the main disulfide reductase, Trx1, is maintained in a reduced state by the action of cytosolic thioredoxin reductase (TrxR1) with potential contribution from both the glutathione reductase (GR) and glutaredoxin (Grx) pathways (Lillig & Holmgren, 2007). Both the Trx1 and glutathione pathways require NADPH as ultimate electron donor to maintain their reductase activity within the cytosol. However, there are no known equivalent pathways present within the ER for the reduction of disulfides in the thioredoxin domains within the PDI family. We recently showed that the regeneration of NADPH within the cytosol is required to ensure correct disulfide formation in the ER lumen (Poet, Oka et al., 2017) raising the possibility that the cytosolic reductive pathways are responsible for ensuring correct disulfide formation in the ER lumen. How the reducing equivalents generated in the cytosol are transferred to the ER lumen remains unknown but a system for the transfer of such equivalents does exist in prokaryotes. Here disulfide formation and reduction within the periplasmic space is catalysed by DsbA and DsbC, which are structurally homologous to Trx1 (Kadokura, Katzen et al., 2003). Disulfide formation is coupled to the electron transport chain via a membrane protein called DsbB, allowing *de novo* disulfide formation in the disulfide exchange protein DsbA. To remove incorrectly oxidised periplasmic thiols, the cytosolic thioredoxin reductase (TrxR) pathway via thioredoxin (Trx) reduces the membrane protein DsbD which transfers electrons across the membrane to reduce DsbC that then catalyses disulfide reduction.

A role for GSH in the reduction of protein thiols has previously been suggested based upon its role as a redox buffer(Chakravarthi, Jessop et al., 2006). This role has been suggested to be required to maintain redox balance after large fluctuations in either reducing or oxidising conditions (Appenzeller-Herzog, Riemer et al., 2010, Jessop & Bulleid, 2004, Molteni, Fassio et al., 2004). A recent report identifying Sec61 as a GSH transporter provides a possible route for its transfer into the ER (Ponsero, Igbaria et al., 2017). However, its requirement for the formation of the correct disulfides in proteins is less clear. Depletion of ER GSH either by inhibition of GSH synthase or by targeting GSH-degrading enzymes does not prevent correct disulfide formation in proteins containing complex disulfides such as tissue-type plasminogen activator or the LDL-receptor (Chakravarthi & Bulleid, 2004, Tsunoda, Avezov et al., 2014). The relative roles of the thioredoxin and GSH pathways in maintaining ER redox poise and reducing oxidised thiols remains an unanswered question (Bulleid & Ellgaard, 2011, Ellgaard et al., 2018).

To evaluate the requirements for the reduction of disulfides within the ER we reconstituted the pathway using purified cytosolic components and microsomal vesicles or semi-permeabilised cells as a source of ER. Using a redox sensitive GFP as a readout (van Lith, Tiwari et al., 2011) we established the minimum requirements for disulfide reduction and demonstrated that the transfer of reducing equivalents across the ER membrane requires a membrane protein. In addition, we show that the resolution of non-native to native disulfides can be driven solely by the thioredoxin pathway. Our results highlight the similarity between the pathways for reduction of disulfides in the bacterial periplasm and the mammalian ER.

## Results

### The reduction of ER-localised disulfides requires an ER membrane component

To follow the reduction of disulfide bonds within the ER lumen, we created a HT1080 stable cell line expressing a version of roGFP that can act as a reporter of disulfide formation within the ER of mammalian cells. To improve the folding and stability of roGFP we included the super-folder mutations as described previously (Hoseki, Oishi et al., 2016), but using an ER targeted roGFP1-iE rather than roGFP1-iL. The resulting cell-line demonstrated bright ER-localised fluorescence that was responsive to changes in both oxidation and reduction making it an ideal reporter for changes in ER redox state (Fig. 1A, B). In addition, there was an absence of light-induced fluorescence changes, an effect that compromised the use of the roGFP1-iL variant (van Lith et al., 2011). The variant of roGFP was designated ER-SFGFP-iE.

**Figure 1:**
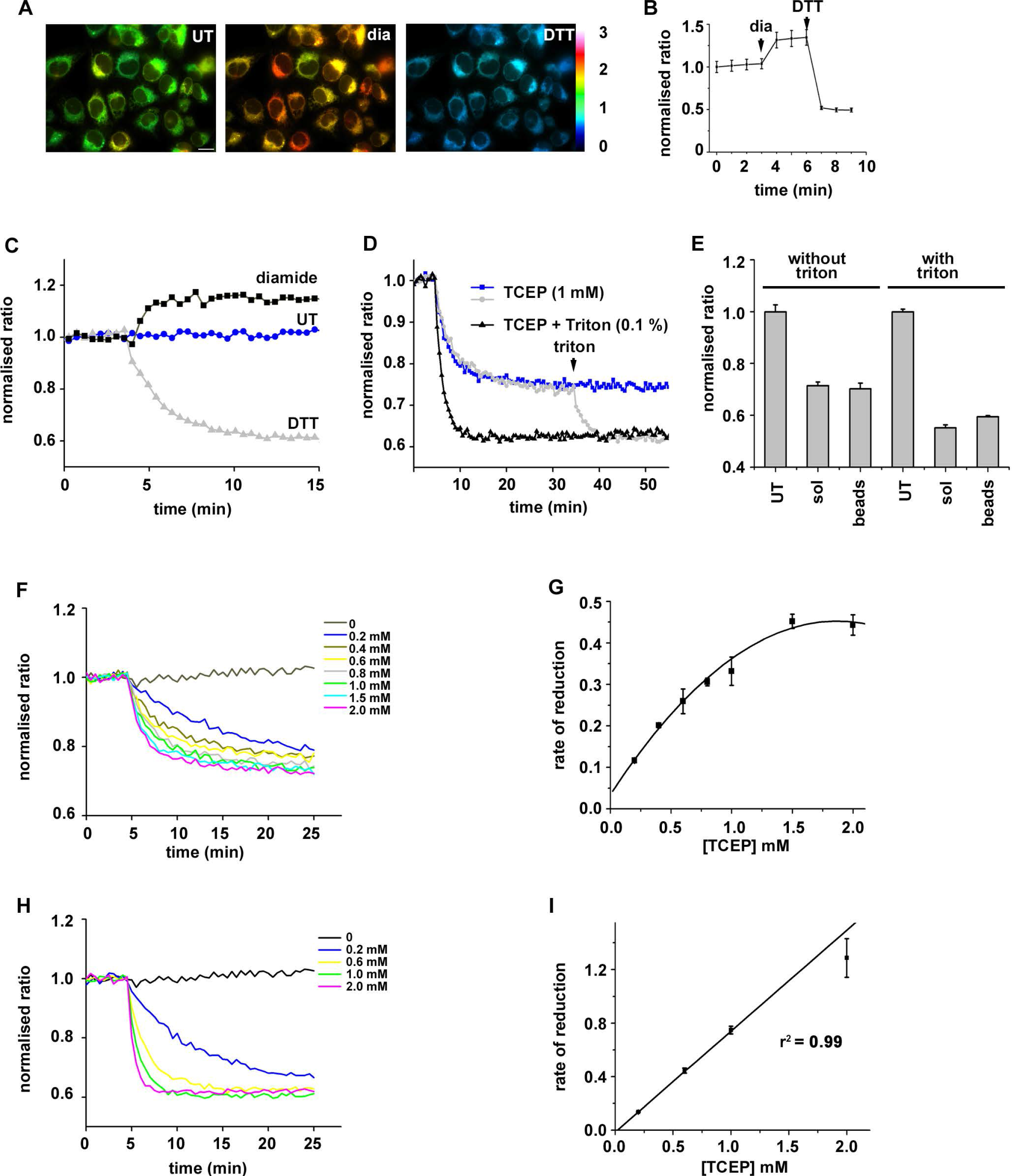
The reduction of ER-localised disulfides by TCEP requires a membrane component. (A and B): A stable cell-line expressing ER-SFGFP-iE was evaluated by live-cell imaging and was responsive to gross changes in redox conditions. (A) Depicts fluorescent images of untreated, diamide (1 mM) or DTT-treated (10 mM) cells. Fluorescence ratio changes are visualised using false colours with a more reducing ratio in blue and a more oxidising in yellow. (B) Normalised fluorescent ratios of individual cells are averaged and changes observed over time by the sequential addition of diamide (1 mM) and DTT (10 mM). (C) Normalised fluorescence ratio changes in microsomal membranes isolated from the ER-SFGFP-iE expressing cell-line followed using a plate reader. An increase in ratio is seen following diamide (1 mM) addition whereas a decrease is seen following DTT (1 mM) addition. The ER-SFGFP-iE in untreated cells is, therefore, neither fully oxidised nor reduced. (D) Changes in normalised fluorescence ratio following the addition of impermeable TCEP in the absence (blue) or presence (black) of Triton X−100 (0.1 % v/v). After 35 min Triton X−100 was added (0.1 % v/v) to solubilise the intact microsomes. Note the slower rate of reduction and the lack of full reduction with TCEP unless the membranes are solubilised. (E) The ER-SFGFP-iE is reduced by TCEP immobilised on agarose beads further confirming that the reduction is not direct. The experiment was carried out in triplicate with average and standard deviation indicated for untreated (UT), soluble (sol) and immobilised (beads) TCEP. Final effective concentration of both soluble and immobilised TCEP was 0.8 mM. (F-I) The change in normalised fluorescent ratio was followed at various TCEP concentrations as indicated either without (F) or with (H) detergent solubilisation of microsomes. The rate of reduction at each TECP concentration was calculated in triplicate with the average +/− standard deviation plotted vs the TCEP concentration (G and I).

We isolated microsomal vesicles from the ER-SFGFP1-iE cell-line and were able to follow changes to ER redox status over time using a plate reader (Fig. 1C). We established that the microsomes were sensitive to both reduction with DTT and oxidation with diamide so the reporter was neither fully oxidised nor reduced following isolation. Reduction with membrane-permeable DTT was rapid reaching completion within 10 min. When the membrane impermeable reducing agent TCEP was added to ER-SFGFP-iE containing microsomes, reduction still occurred but much slower than if the microsomes were solubilised priot to TCEP addition (Fig. 1D, blue vs black). Reduction in intact microsomes did not reach completion unless detergent was added to solubilise the ER membrane (see from 35 min). To ensure the TCEP was not directly reducing ER-SFGFP-iE, we carried out a similar experiment but used TCEP immobilised on agarose beads. ER-SFGFP-iE was reduced by either immobilised or free TCEP to an equivalent ratio with full reduction only occurring following membrane solubilisation. These results strongly suggest the indirect reduction of ER-SFGFP-iE by TCEP when the microsomal membrane is intact.

The lack of full reduction of ER-SFGFP-iE by TCEP suggests that the reaction had reached saturation. To evaluate this possibility further we determined the rate of reduction with a range of TCEP concentrations (Fig. 1F, G). The results show that the rate of reduction does indeed reach saturation at concentrations of TCEP over 1 mM suggesting a facilitated process. To show that TCEP can reduce ER-SFGFP-iE directly and fully we determined the rate of reduction with a range of TCEP concentrations after solubilising the ER membrane (Fig. 1H, I). Under these conditions the rate did not reach saturation suggesting a diffusion limited process. These results with the membrane impermeable reducing agent suggest the indirect reduction of an ER-localised disulfide mediated via an ER membrane component.

### The thioredoxin reduction pathway is sufficient to reduce ER localised proteins via a membrane protein

To evaluate the cytosolic components required to ensure reduction of ER-localised disulfides, we attempted to reconstitute the pathway with purified components. We reasoned that a source of NADPH would be required as well as TrxR1 and Trx1. To ensure efficient recycling of NADPH we included glucose 6 phosphate (G6P) as well as G6P dehydrogenase. To demonstrate that the system could maintain Trx1 in its reduced state we determined the redox status of purified Trx1 before and after incubation with the recycling system. Redox status was evaluated following modification with 4-acetamido-4’-malemidylstilbene-2,2’-disulfonic acid (AMS) which increases the protein mass slowing its electrophoretic mobility. Prior to incubation, Trx1 was present as a mixture of both reduced and oxidised protein (Fig. 2A, lane 1). After incubation with TrxR1 and the NADPH recycling system Trx1 was fully reduced (lane 2). When our ER-SFGFP-iE containing microsomes were incubated with the TrxR1 reduction system, roGFP became more reduced, an effect that was dependent upon the inclusion of Trx1 as no reduction occurred in the presence of TrxR1 and the NADPH recycling system alone (Fig. 2B, C-Trx)). No reduction of roGFP was observed when active site mutants of Trx1 (CXXS or SXXS) were used in place of wild type protein (CXXC). These results demonstrate that the minimal requirement for reduction of ER-localised roGFP is the thioredoxin reduction system and that disulfide exchange is mediated though the Trx1 CXXC active site motif. As the reduction was Trx1-dependent which is membrane impermeable, it is highly likely that an ER membrane component is required to transfer the reducing equivalents from Trx1 to roGFP.

**Figure 2:**
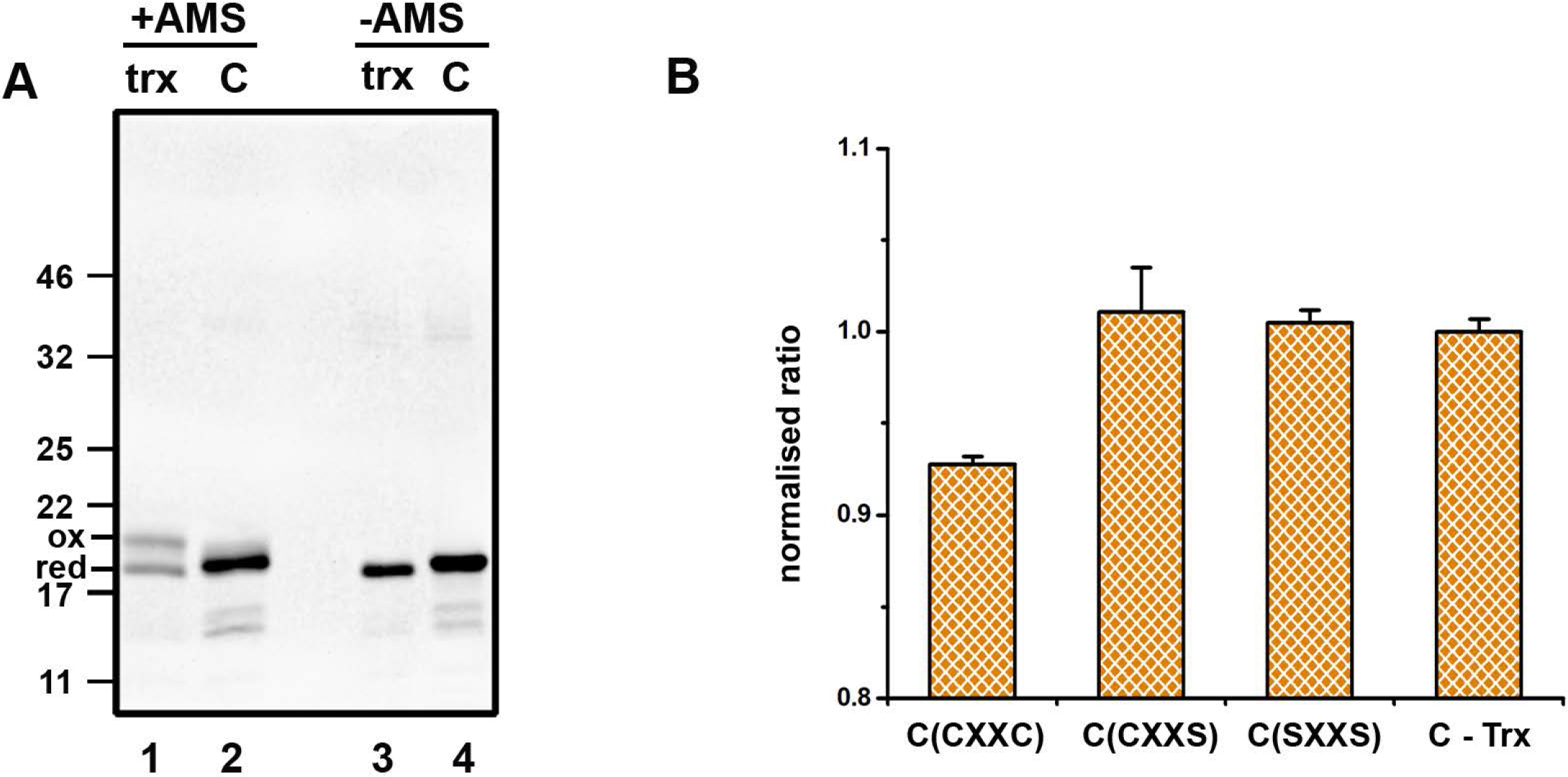
Thioredoxin-dependent reduction of ER-localised disulfides. (A) The redox status of Trx1 (trx) was determined by AMS differential alkylation either before (lane 1) or after (lane 2) incubation with TrxR1 and a NADPH recycling system (C). None AMS-treated samples are shown in lanes 3 and 4. (B) The normalised fluorescence ratio of ER-SFGFP-iE was measured following incubation with the reduction system without Trx1 (C-Trx) or with the TrxR1 system containing wild type Trx1 (CXXC) or active site mutants CXXS or SXXS. The experiment was repeated three times with the average and standard deviation of the three replicates presented.

The experiments with roGFP provide us with a useful way to follow reduction of a specific disulfide within the ER lumen by components added outside the vesicles, however, this protein is not normally present in the ER. Hence, we evaluated the redox status of ERp57, a member of the PDI family, to determine whether the thioredoxin reduction system could also reduce an endogenous ER protein that has previously been shown to be required for the correct folding of glycoproteins (Jessop et al., 2007). We used an approach based upon the sequential alkylation of cysteine residues (Jessop & Bulleid, 2004) to probe the redox status of ERp57 in semi-permeabilised (SP)-cells isolated from a cell-line stably transfected with V5-tagged ERp57 (Jessop et al., 2007). Samples were either untreated, treated with the oxidising agent diamide, with the reducing agent DTT or with the non-membrane permeable TCEP (Fig. 3A). Treatment with NEM blocks any free thiols and subsequent reduction and alkylation with AMS reduces the mobility of the protein. Proteins that contain disulfides in the original SP-cells will, therefore, have reduced mobility on SDS-PAGE. Following this procedure, ERp57 in untreated SP-cells migrated as two species representing the oxidised and reduced forms by comparison with the diamide or DTT treated samples (lanes 1-3). After incubation with TCEP most of the ER-localised ERp57 was reduced though not completely (lane 4), a result which mirrors the incomplete reduction of roGFP and demonstrates that TCEP can substitute for a cytosolic reduction system.

**Figure 3:**
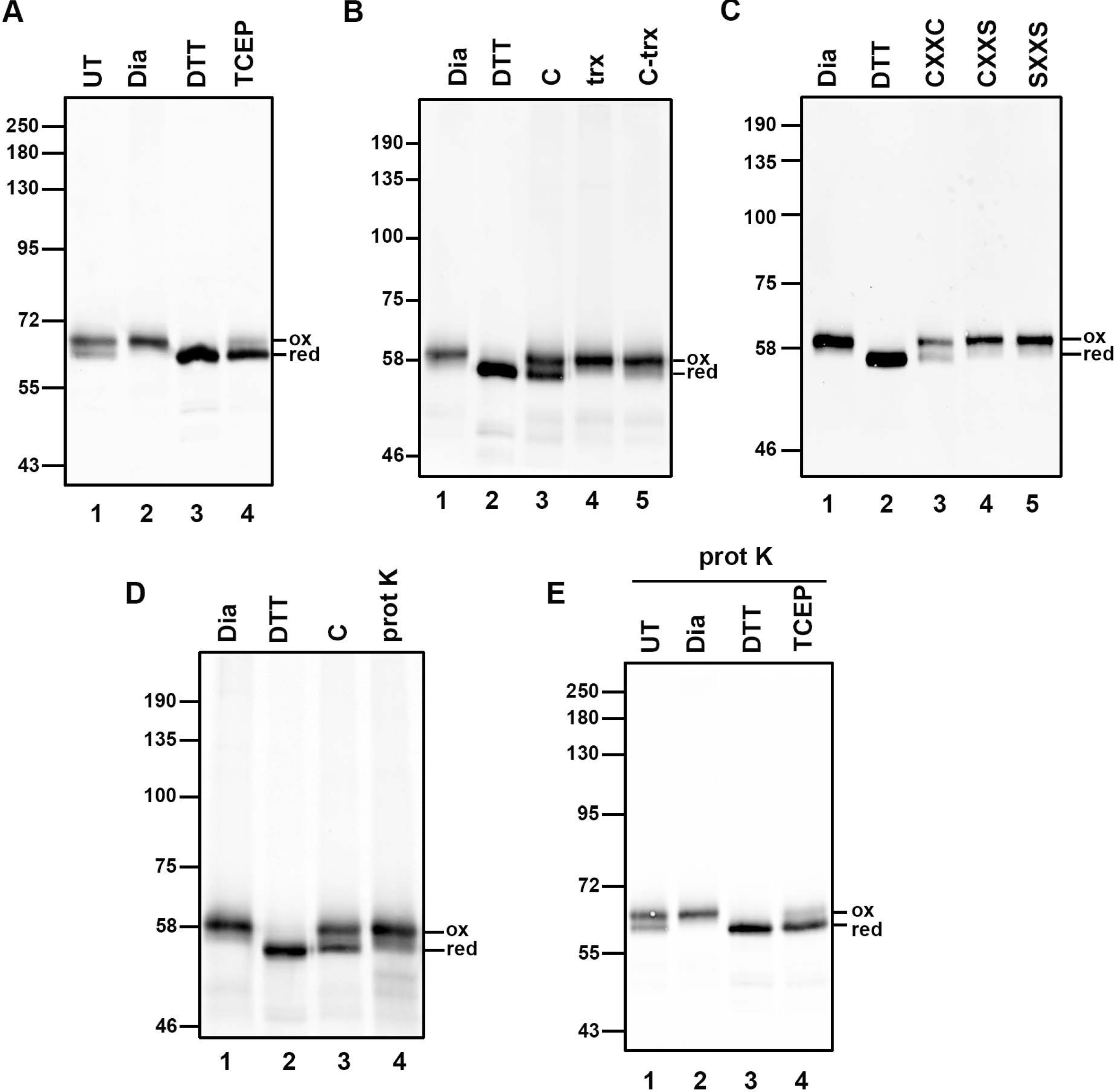
Thioredoxin-dependent reduction of ERp57 requires a membrane protein. (A) The redox status of ER-localised ERp57 was determined following differential alkylation with AMS. In untreated SP-cells, ERp57 exists as a mixture of reduced and oxidised forms (lane 1) and becomes fully oxidised (lane 2) or fully reduced (lane 3) following the addition of diamide or DTT respectively. Partial reduction of ERp57 also occurs following incubation of fully oxidised ERp57 with the membrane impermeable reductant TCEP (lane 4). (B) Thioredoxin-dependent reduction of ER-localised ERp57. SP-cells were treated with diamide before incubation with either the complete TrxR1 reduction system with (C - lane 3) or without (C – Trx, lane 5) or with Trx1 alone (lane 4). ERp57 was partially reduced only by the complete system. (C) The reduction of ERp57 by the TrxR1 system needs Trx1 with an intact active site. Neither of the active site mutants CXXS (lane 4) or SXXS (lane 5) can substitute for wild type (CXXC) Trx1. (D) The reduction of ER-localised ERp57 by the TrxR1 system is sensitive to prior treatment of the SP-cells with proteinase K (lane 4). (E) The reduction of ER-localised ERp57 by TCEP is not prevented by prior treatment of the SP-cells with proteinase K (lane 4).

To evaluate the ability of the thioredoxin pathway to reduce ER-localised ERp57, we first fully oxidised SP-cells with diamide, removed diamide by re-isolating the SP-cells and incubated with either the complete TrxR1 reduction system, with just recombinant Trx1 or with the reduction system without Trx1 (Fig. 3B). Partial reduction of fully oxidised ERp57 occurred in the presence of the complete system (lane 3) but did not occur with just Trx1 alone (lane 4) indicating the requirement for the recycling system. The reduction of ERp57 was dependent upon Trx1 (lane 5) as was the case with roGFP. In addition, the active site mutants of Trx1 lacking one or both cysteines within the CXXC motif were unable to reduce ERp57 (Fig. 3C, lanes 4 and 5). These results demonstrate that ER-localised ERp57 can be reduced following oxidation by the cytosolic reduction pathway and that this reduction is dependent upon thioredoxin.

To further characterise the membrane component required for reduction, we carried out a proteinase K digestion of our SP-cells prior to assaying for ERp57 reduction (Fig. 3D). We surmised that if a membrane protein was involved then the transfer of reducing equivalents across the membrane may be sensitive to proteolysis. Proteinase K digestion prevented the reduction of oxidised ERp57 by the reconstituted reduction system (lane 4) suggesting the requirement for a membrane protein whose cytosolic domain is sensitive to digestion. We also determined whether the reduction of ERp57 by TCEP was sensitive to proteinase K treatment of the SP-cells (Fig. 3E). TCEP was still able to reduce oxidised ERp57 (lane 4) suggesting that either the membrane protein involved can be reduced efficiently by TCEP without its protease sensitive cytosolic domain or that TCEP acts via a separate component than the thioredoxin-dependent pathway.

### The thioredoxin reduction pathway is sufficient to reduce non-native disulfides in nascent chains exposed to the ER lumen

We previously demonstrated that non-native disulfides formed during the synthesis of β1-integrin require a cytosolic reductive pathway to ensure their isomerisation to the correct disulfides (Poet et al., 2017). To evaluate the ability of our reconstituted pathway to isomerise non-native disulfides, we generated stalled translocation intermediates of a protein that forms non-native disulfides following *in vitro* translation in the presence of semi-permeabilised (SP) cells (Robinson, 2019). By isolating SP-cells containing these intermediates we were then able to follow the rearrangement of disulfides post-translationally in the presence or absence of a TrxR1 reduction system (Fig. 4A). We generated a stalled translocation intermediate of disintegrin domain of ADAM10 by translating an RNA transcript lacking a stop codon. The length of the synthesised chain is sufficient to allow exposure of several cysteines in the ER lumen (Fig. 4B) that have the potential to form disulfides. Translations were performed using a rabbit reticulocyte lysate supplemented with SP-cells to study folding in the ER (Wilson, Allen et al., 1995). All samples were treated with NEM on completion to irreversibly modify thiols and freeze the disulfide status of the samples for downstream processing. When translations were carried out in the presence of a reducing agent (DTT) and the samples separated under non-reducing conditions, two products of approximately 18 and 20 kDa were synthesised (Fig. 4C, lane 1). These products correspond to the translocated, signal sequence cleaved chain and untranslocated preprotein (Robinson, 2019). When the translations were carried out in the absence of added G6P and a post-translational incubation carried out in the absence of G6P, a diffuse pattern emerged with the translation products forming both intra and inter-chain disulfides (lane 2). When G6P was added post-translationally and samples incubated for 60 min, more distinct products were formed indicative of discrete disulfide-bonded species (lane 3). When SP-cells were isolated from translations carried out in the absence of G6P and incubated in KHM buffer alone, no change to the banding pattern was observed (lane 4). However, when the TrxR1 reduction system was added to the isolated SP-cells, rearrangement of disulfides was observed resulting in more distinct disulfide-bonded species (lane 5). This change to the banding pattern to more discrete disulfide-bonded species did not occur when the isolated SP-cells were incubated in the TrxR1 reductive pathway lacking Trx1 (lane 6). We also demonstrated that TCEP could substitute for G6P when added to translations reactions post-translationally and to isolated SP-cells post-translational giving rise to discrete disulfide-bonded species (Fig. 4C, lanes 3 and 5). We have previously shown that the disulfide-bonded products formed form this ADAM10 construct are present in nascent chains translocated into the ER lumen (Robinson, 2019). Taken together these results show that a source of non-membrane permeable reducing agent can resolve the non-native disulfides formed in nascent chains within the ER lumen.

**Figure 4:**
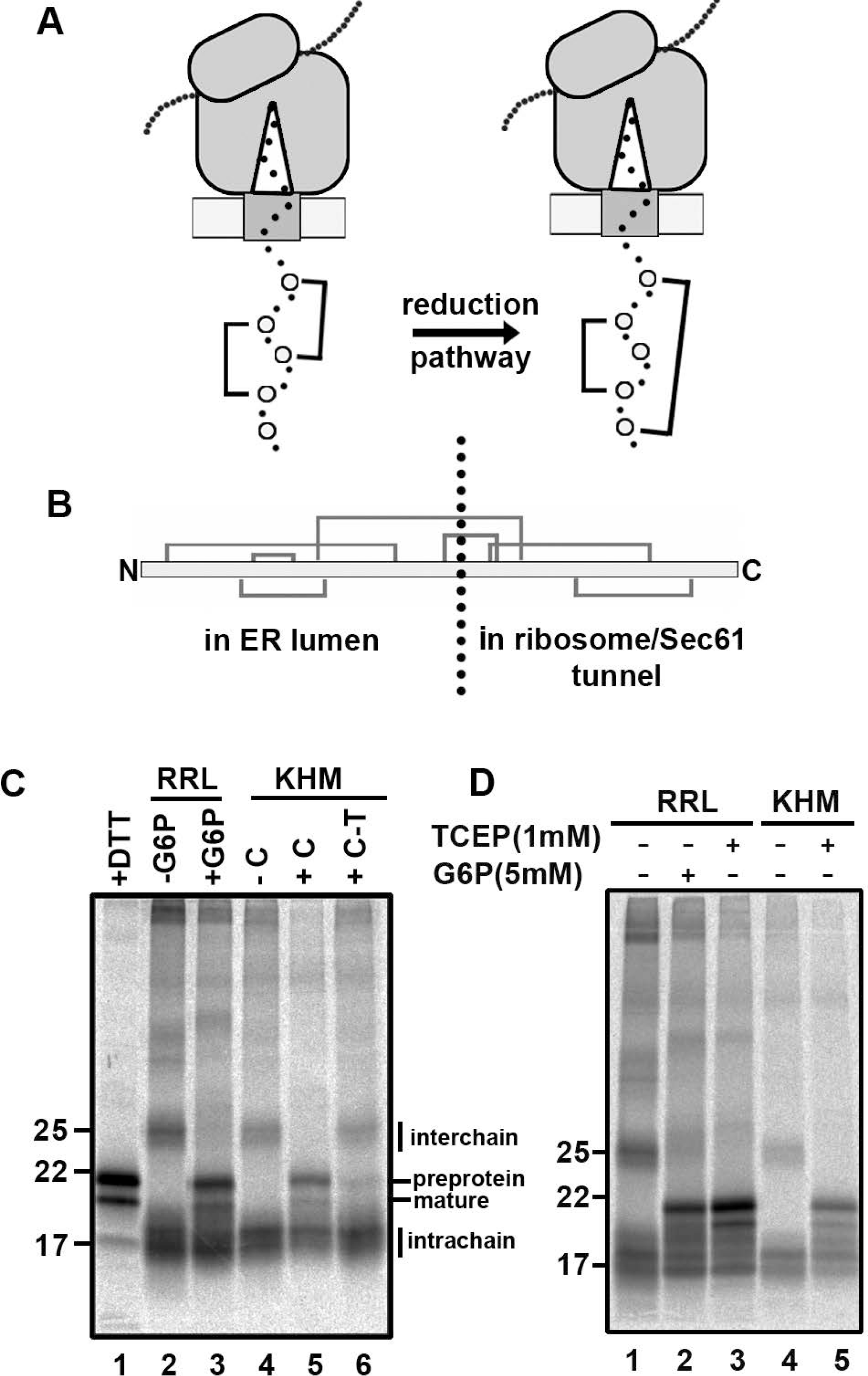
Thioredoxin-dependent post-translational rearrangement of ER-localised nascent chain disulfides. (A) Cartoon depicting the rearrangement of nascent chain disulfides in a stalled translocation intermediate following the addition of a cytosol-localised reduction system. (B) Schematic of the disulfide pattern within the disintegrin domain of ADAM10. The stalled intermediate used in subsequent experiments will have the indicated disulfide-forming cysteines in the ER lumen. (C) Translations carried out in the presence of DTT (5 mM) gave rise to distinct translation products which migrate with the sizes of the pre and mature protein as indicated (lane 1). Translation were carried out in the absence of added G6P and SP-cells subsequently incubated post-translationally in a reticulocyte lysate (RRL) in the absence (lane 2) or presence (lane 3) of G6P. SP-cells were isolated from the translations and resuspended in KHM buffer alone (- C) (lane 4). Note the rearrangement of nascent chain disulfides when the TrxR1 system is included in the incubation (lane 5) (+ C) and the dependence of this rearrangement on the presence of Trx1 (lane 6) (C − T). (D) Similarly to the TrxR1 system, TCEP brings about the rearrangement of nascent chain disulfides when added to SP-cells post-translationally either in the presence of the rabbit reticulocyte lysate (lane 3) or in KHM buffer alone (lane 5).

### The thioredoxin reduction pathway reduces disulfides in several secretory and ER resident proteins

The experiments with roGFP and ERp57 are dependent upon the reduction of a reversible disulfide that is solvent accessible in the final protein structure. To determine whether our reconstituted reductive system could reduce other disulfides within secretory or membrane proteins, we carried out redox proteomics. Essentially microsomes were incubated with the TrxR1 system in either the presence or absence of Trx1 for 60 min and reversibly oxidised thiol groups were labelled with heavy-iodoacetamide. The samples were prepared for LC/MS/MS analysis by trypsin digestion followed by tandem mass tagging to enable quantification of the changes in the cysteine oxidation levels between experimental conditions. The resulting data was analysed by Perseus software and is presented as a volcano plot indicating the statistically significant changes to the reversibly oxidised cysteine residues upon incubation with Trx1 (Fig. 5A). The redox status of several peptides changed during the incubation, specifically there was a subset of cysteines whose oxidised status decreased (more reduced) in the reactions containing Trx1, with the majority of the corresponding proteins being located in the ER lumen (blue circles). Notably the regulatory cysteine (131) in Ero1α was reduced (Appenzeller-Herzog, Riemer et al., 2008, Baker, Chakravarthi et al., 2008) as was the cysteine involved in recycling VKORC1L1 (50) (Tie, Jin et al., 2014). Changes to the redox status of a few cytosolic and mitochondrial proteins also occurred (orange circles), which likely co-purified with our microsomal vesicles. The identity of all the proteins identified whose redox status changed along with their ultimate subcellular location is as illustrated (Fig. 5B) or detailed separately (Table 1). The results demonstrate that several thiols within endogenous proteins that are synthesised at the ER, including thiols that form structural and regulatory disulfides, can be reduced by the thioredoxin reduction system. The target proteins are localised to the ER lumen and the TrxR1 reduction system is membrane impermeable illustrating the requirement to transfer reducing equivalents across the membrane.

**Figure 5:**
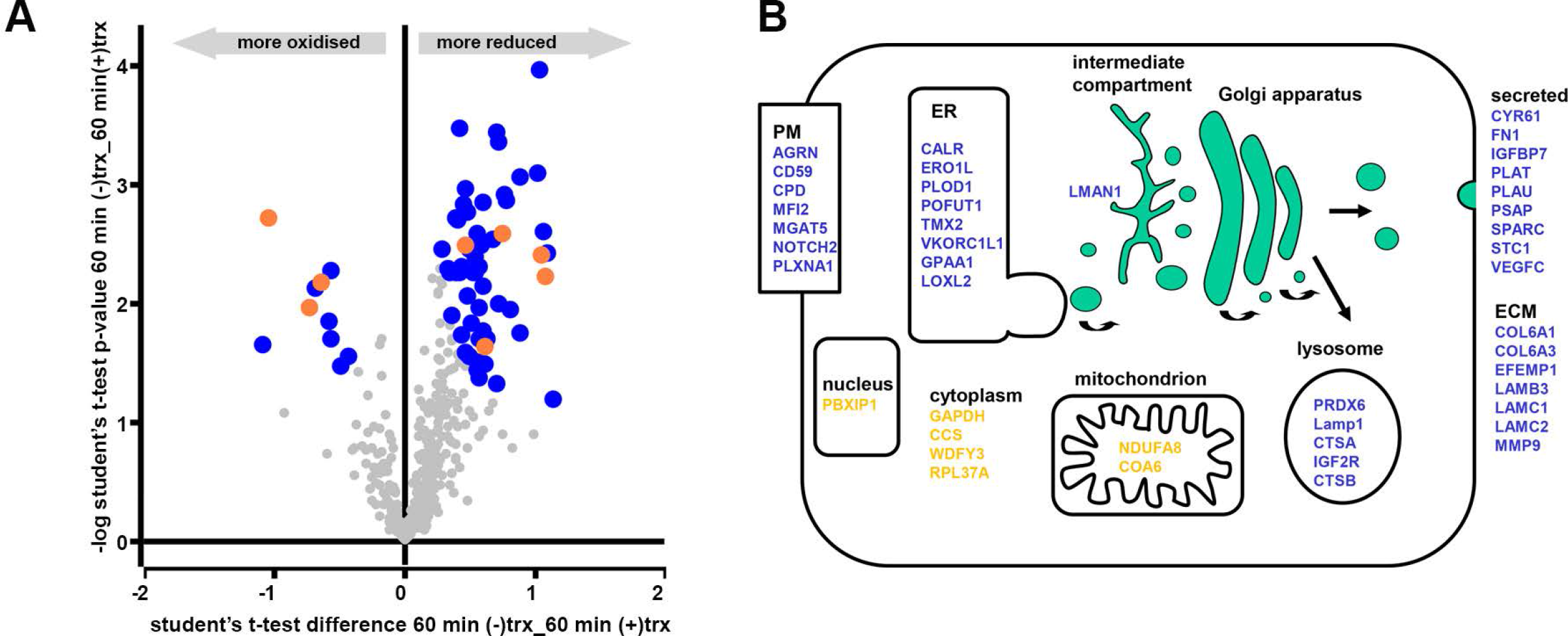
Evaluation of the changes in the redox status of microsomal proteins following incubation with the TrxR1 system. (A) Volcano plot depicting proteins with significant changes to their redox status where the modified cysteines is localised to the ER (blue) or the cytosol (orange). Proteins identified by mass spectrometry whose redox status does not change significantly are also indicated (grey). (B) Cartoon illustrating the final cellular location of proteins whose redox status changes significantly upon incubation of microsomes with the TrxR1 system. Proteins that enter the secretory pathway are in blue whereas those that do not enter the ER are in orange.

**Table 1:** List of proteins showing a significant change to their oxidised thiols along with information on their subcellular location, position and identity of the modified cysteine(s).

| <u>Location of cys modified</u> | <u>Gene name</u> | <u>protein location</u> | <u>type</u> | <u>Cysteine modified</u> |
| --- | --- | --- | --- | --- |
| **cytosol** | GAPDH | cytoplasm | soluble | 152;156 |
|  | CCS | cytoplasm | soluble | 227 |
|  | WDFY3 | cytoplasm | soluble | 952 |
|  | RPL37A | cytoplasm | soluble | 39;42 |
|  | KIDINS220 | endosome | multi-pass | 705 |
|  | PBXIP1 | nucleus | soluble | 559;565 |
| **ER** | COL6A1 | ECM | soluble | 745 |
|  | COL6A3 | ECM | soluble | 1823;1824 |
|  | EFEMP1 | ECM | soluble | 40;47 |
|  | LAMB3 | ECM | soluble | 451;453;511;534;536 |
|  | LAMC1 | ECM | soluble | 416;418;474;751;804;807;954; 956 |
|  | LAMC2 | ECM | soluble | 28;30;53;56 |
|  | MMP9 | ECM | soluble | 256 |
|  | CALR | ER resident | soluble | 105 |
|  | ERO1L | ER resident | soluble | 131 |
|  | PLOD1 | ER resident | soluble | 267;270 |
|  | POFUT1 | ER resident | soluble | 283 |
|  | TMX2 | ER resident | single-pass | 59 |
|  | GPAA1 | ER resident | multi-pass | 266 |
|  | VKORC1L1 | ER resident | multi-pass | 50 |
|  | LMAN1 | ER/Golgi intermediate compartment | single-pass | 230 |
|  | LOXL2 | ER/secreted | soluble | 395 |
|  | PRDX6 | lysosomal | soluble | 47 |
|  | LAMP1 | lysosomal | single-pass | 338 |
|  | CTSA | lysosomal | soluble | 239 |
|  | IGF2R | lysosomal | soluble | 1227 |
|  | CTSB | lysosome/secreted | soluble | 211 |
|  | CD59 | plasma membrane | GPI-anchored | 70 |
|  | CPD | plasma membrane | single-pass | 309 |
|  | MFI2 | plasma membrane | GPI-anchored | 199 |
|  | MGAT5 | plasma membrane | single-pass | 338 |
|  | NOTCH2 | plasma membrane | single-pass | 1342;1346 |
|  | PLXNA1 | plasma membrane | single-pass | 358 |
|  | AGRN | secreted | soluble | 650 |
|  | CYR61 | secreted | soluble | 337;314;317 |
|  | ENDOD1 | secreted | soluble | 34 |
|  | FN1 | secreted | soluble | 87;97;135;141;179;231;508 |
|  | IGFBP7 | secreted | soluble | 156 |
|  | PLAT | secreted | soluble | 419 |
|  | PLAU | secreted | soluble | 45 |
|  | PSAP | secreted | soluble | 42 |
|  | SPARC | secreted | soluble | 130 |
|  | STC1 | secreted | soluble | 170 |
|  | VEGFC | secreted | soluble | 328 |
| **mitochondrion** | NDUFA8 | mitochondrion | soluble | 66 |
|  | COA6 | mitochondrion | soluble | 44 |
| **unknown** | NUMB | peripheral membrane protein | unknown | 130;132 |
|  | DPY19L1 | Unknown | multi-pass | 647 |

## Discussion

Our previous work provided the first indication that a cytosolic reductive pathway was required to ensure correct disulfide formation within the ER (Poet et al., 2017). Here we demonstrate that the thioredoxin pathway is sufficient to provide the reducing equivalents to ensure the reduction of regulatory and non-native disulfides within ER proteins. We have successfully reconstituted the reduction of ER disulfides using a microsomal system in the presence of purified components to demonstrate that the thioredoxin pathway can drive the reduction of disulfides in several ER-localised soluble and membrane proteins. In addition, we show that the pathway is sufficient to facilitate the reduction of disulfides formed in nascent chains entering the ER lumen but does not prevent disulfide reformation – a requirement for correct isomerisation of non-native to native disulfides. Importantly we show that a membrane protein is required to facilitate the transfer of reductant across the ER membrane, presumably via a disulfide exchange mechanism as demonstrated for the transfer of disulfides across the bacterial plasma membrane by DsbD (Cho, Porat et al., 2007).

The absolute requirement for the presence of thioredoxin in our reconstituted system rules out a role for an NADPH-dependent ER-localised reductase. Previously it has been postulated that an ER glutathione or thioredoxin reductase could utilise the ER pool of NADPH to drive the reduction of disulfides (Bulleid & Ellgaard, 2011). As there is an ER-localised hexose 6-phosphate dehydrogenase (Lavery, Walker et al., 2006), G6P may enter the ER via the G6P-transporter, be metabolised to phosphogluconolactone and in the process regenerate the ER pool of NADPH. Hence, the G6P effect we previously observed during our *in vitro* translations may be due to a process independent of the thioredoxin pathway. This is clearly not the case for the reduction of roGFP, ERp57, disulfides in nascent chains and the range of proteins identified from the redox proteomics whose redox state changed depended upon the presence of thioredoxin.

The reductive pathway driven by cytosolic thioredoxin is versatile in terms of its target disulfides in the ER lumen. These disulfides included those present within folded proteins such as roGFP, in proteins undergoing folding during translocation and in proteins that require reduction to maintain their activity. Such versatility would suggest a broad substrate specificity of the terminal enzyme involved in catalysing reduction. Changes to the redox status of ERp57 would suggest that this member of the PDI family could act as a reductase as suggested previously for newly synthesised glycoproteins (Jessop et al., 2007) to which it is targeted via its interaction with calnexin or calreticulin (Oliver et al., 1997). It is also likely that ERdj5 is reduced by the cytosolic reductive pathway and be targeted to substrate proteins through its interaction with BiP (Cunnea et al., 2003). Our identification of one of the regulatory disulfides within Ero1α (Baker et al., 2008) as a target for the cytosolic reductive pathway would suggest that this disulfide can be reduced by one of the PDI family members as demonstrated previously using purified proteins (Shepherd, Oka et al., 2014). Whether the ER membrane protein responsible for transfer of reducing equivalents directly reduces the PDI reductase remains to be established.

The role of thioredoxin as reductant of the ER membrane protein can be replaced with the non-membrane permeable reducing agent TCEP. However, in contrast to thioredoxin-mediated process, reduction of ER proteins via TCEP was shown to be insensitive to protease digestion suggesting that this reagent can reduce the ER membrane protein in the absence of a protease sensitive cytosolic domain. This result most likely reflects the different requirements for reduction of the membrane protein disulfide by thioredoxin and TCEP. The mechanism underlying recognition and reduction of substrates by thioredoxin is not completely understood given its wide range of substrate specificity. However, it has been demonstrated that non-covalent interactions prior to disulfide reduction are important with recognition being due to the conformational restriction afforded by the oxidised substrate (Palde & Carroll, 2015). Hence the proteolytic digestion of a protein domain or loop may prevent thioredoxin interacting with the membrane protein thereby inhibiting activity. A chemical reductant such as TCEP would have no requirement for such non-covalent interactions and would only require the target disulfide to be solvent accessible.

The identity of the membrane protein involved connecting the cytosolic thioredoxin couple with the ER remains unknown. One resident ER membrane protein whose depletion leads to a more oxidising ER and a specific defect in correct disulfide formation in lipoprotein lipase is lipase maturation factor 1 (LMF1) (Roberts, Babilonia-Rosa et al., 2018). This protein contains several thiols in both its cytosolic loops, transmembrane regions as well as its lumenal domain that are required for activity and could be involved in disulfide exchange. Unfortunately, we did not identify LMF1 in our proteomics data, so we could not identify any changes to its redox status. Hence, LMF1 remains a potential candidate to shuttle disulfides across the ER membrane. Of the 41 microsomal proteins whose redox status changes during the incubation with the TrxR1 system, 4 are multi-pass membrane proteins with VKORC1L1 and GPAA1 being known resident ER proteins. GPAA1 is a subunit of the GPI-anchor transamidase (Ohishi, Inoue et al., 2000) whereas VKORC1L1 is a paralogue of vitamin K epoxide hydrolase with a potential function in vitamin K-dependent carboxylation (Tie, Jin et al., 2013). Whether these proteins are directly involved in disulfide exchange across the ER membrane is a focus of future studies.

## Materials and Methods

### Generation of stable cell line expressing ER-SFGFP-iE

HT1080 cells were transfected with ER-SFGFP-iE using MegaTran 1.0 (OriGene, Cambridge, UK) following the manufacturer’s protocol. Various dilutions (1:10 to 1:10,000) of transfected HT1080 cells were plated on 15 cm^2^ dishes. Cells were grown in selective medium containing G418 (1mg/ml) until colonies appeared. Single colonies were picked and transferred to 12 well plate (one colony per well) using trypsin-soaked Scienceware^®^ cloning discs (Sigma. Dorset, UK). Cells were grown until confluent and then transferred to 25 cm^2^ flasks. The presence of ER-SFGFP-iE was confirmed by western blot using anti-GFP antibody (Santa Cruz, California, USA).

### Live cell microscopy and image analysis

HT1080 cells stably expressing ER-SFGFP-iE were seeded on coverslips. The cells were washed with buffer (HEPES (20 mM) pH 7.4, NaCl (130 mM), KCl (5 mM), CaCl_2_ (1 mM), MgCl_2_ (1 mM), D-glucose (10 mM)) before imaging on a Zeiss Axio Observer A1 inverted microscope equipped with a 40 x FLUAR lens (Zeiss, Oberkochon, Germany). The cells were sequentially excited by 385 and 470 nm LEDs with fluorescence detection through a T495 lpxr beamsplitter (Chroma, Vermont, USA). Images were captured at 30 s intervals and analysed with AxioVision 4.8 software. The 385/470 nm ratio images of untreated, diamide-treated (1 mM) and dithiothrietol (DTT)-treated (10 mM) cells were false-coloured using a rainbow look-up table. The 385/470 nm ratio changes over time were quantified by defining regions of interest for individual cells and plotted as the average ratio with standard deviation.

### Preparation of microsomal membranes

Cells were cultured using cell culture roller bottles (1700 cm^2^) (Greiner Bio-one). Confluent cells were detached from the roller bottle by incubation with 100 ml PBS containing EDTA (1 mM) for 15 min. Cells were pelleted, washed three times in HEPES–KOH (35 mM) buffer pH 7.5, containing NaCl (140 mM) and glucose (11 mM) followed by washing once in extraction buffer (HEPES–KOH (20 mM) pH 7.5, KOAc (135 mM), KCl (30 mM), and MgOAc(1.65 mM)). An equal volume of extraction buffer to the cell volume was used for resuspension. The cells were disrupted in a Mini-Bomb Cell disruption chamber (KONTES) using a nitrogen pressure of 1.0 MPa for 30 min. Cell homogenates were centrifuged at 6000 × g for 5 min at 4 °C and the post-nuclear supernatant was retained. The supernatant was centrifuged at 50,000 × g (Optima™ Max-XP Ultracentrifuge, Beckman Coulter) for 15 min at 4 °C to pellet the membrane fraction. The membrane pellet was resuspended in buffer A (Tris-HCl (50 mM) pH7.4, sucrose (0.25 M), KCl (50 mM), MgOAc (6 mM), EDTA (1 mM)) to give an A_280_ value of 50 units/μl (the absorbance was determined in water in the presence of 1% SDS). Aliquots of the microsomal membranes were frozen in liquid nitrogen and stored at −80 °C.

### Measurement of changes to the redox state of microsomal ER-SFGFP-iE

Microsomes were pipetted into Dulbecco's phosphate-buffered saline (PBS) (1.5 μl microsomes in 48.5 μl PBS) in the absence or presence of 0.1% Triton X-100 (Thermo Fisher). The solution was added into a CELLSTAR™ 96 Well Polystyrene Flat Bottom Cell Culture Microplate (Greiner Bio-One) (final volume 50 μl/well) (three wells per condition). Fluorescence intensities at 520 nm were measured following excitation at 390nm and 460 nm excitations wavelengths using PHERAstar^®^ FS microplate reader (BMG LABTECH). After 10 min measurement to obtain a baseline, DTT, diamide or different concentrations of Tris-(2-Carboxyethyl)phosphine (TCEP) (Thermo) were injected into the wells. Fluorescence intensities were recorded for up to 50 min. In some experiments, the same amount of microsomes were untreated or treated with DTT, diamide or immobilized TCEP Disulfide Reducing Gel (Thermo) which has been washed twice with PBS to remove soluble TCEP. The samples were then transferred into microplate wells to measure fluorescence intensity at 520 nm following excitation at 390 and 460 nm excitation.

To calculate the rate of change of the redox status of ER-SFGFP-iE at different TCEP concentrations, microsomes were treated with varied concentrations of TCEP in absence or presence of Triton X−100 (0.1% v/v). The fluorescence ratio (390nm/460nm) change was followed over time after addition of TCEP and the resulting time course fitted to an exponential decay function (y = −A(e^−k(t−t0)^−1) + c) to calculate the rate (k), where t_0_ = time of addition of TCEP, c = initial fluorescence ratio and A = difference in fluorescence ratio between t_0_ and t. The rate was then plotted against the TCEP concentration. The rate of reduction of ER-SFGFP-iE at different concentrations of TCEP was performed in triplicate.

### Recombinant protein purification

Recombinant human thioredoxin (Trx1) wild-type and mutants were expressed in and purified from *Escherichia coli* BL21-DE3 cells as described previously (Schwertassek, Balmer et al., 2007). Purification was by successive HisTrap affinity and size exclusion (Superdex 200 10/300) chromatography (GE Healthcare). The protein concentration was determined following OD_280nm_ measurement and calculated using an extinction coefficient of 12.49 mM^−1^cm^−1^. The purified protein aliquots were flash frozen in liquid nitrogen and then stored −80 °C.

### Determining the redox state of Trx1

Purified Trx (25 μM) was incubated in KHM buffer (HEPES (20 mM) buffer pH 7.2, including KOAc (110 mM), MgOAc (2 mM)) at 30 °C for 60 min in the absence or presence of NADPH (1 mM), human TrxR1 (16.25 nM) (IMCO), G6P (1.25 mM) and G6PDH (10 U/ml). The reaction was stopped by the addition of NEM (20 mM). Samples were incubated with streptavidin agarose resin (Thermo) in isolation buffer (Tris-HCl (50 mM) pH7.5, Triton (1%), NaCl (150 mM), EDTA (2 mM), PMSF (0.5 mM) and Na-azide (0.02%w/v)) at 4 °C for 60 min. The streptavidin beads were pelleted by centrifugation and washed three times in isolation buffer to remove unbounded proteins. Trx1 was eluted from the streptavidin beads by incubating at room temperature for 15 min with PBS including SDS (2 % w/v) and biotin (3 mM). The samples were boiled at 105 °C for 15 min and incubated with TCEP (10 mM) at room temperature for 10 min to break existing disulfides followed by a 1.5 h incubation with AMS (20 mM) (Invitrogen) in the dark to alkylate free thiols. The samples were separated by SDS-PAGE and a Western blot performed using Streptavidin Protein DyLight 800 (ThermoFisher).

### Preparation of semi-permeabilised (SP) cells

SP-cells were prepared as described previously (Wilson, Allen et al., 1995). Following digitonin treatment, cells were resuspended in KHM buffer containing CaCl_2_ (1 mM) and treated with *Stapylocossus aureus* nuclease (Calbiochem) (150 U/ml) at room temperature for 12 min to remove the endogenous mRNA. Ethylene glycol-bis(2-aminoethylether)-N,N,N′,N′-tetraacetic acid (EGTA, Sigma) (4.5 mM) was used to stop the reaction. SP-cells were pelleted by centrifugation and resuspended in 100 μl KHM buffer to be used in translation reactions. For ERp57 redox state determination experiments, SP-cells were prepared from cells expressing V5-tagged ERp57 (Jessop et al., 2007) and nuclease treatment was not included during the preparation.

### Determining the redox state of ER-localised ERp57

The redox state of ERp57 in SP-cells was determined as described previously (Jessop & Bulleid, 2004). SP-cells were either untreated or treated with diamide (10 mM) at room temperature for 10 min. Samples were washed twice with 500 μl KHM buffer to remove diamide. Oxidised SP cells was incubated at 30 °C for 60 min in KHM buffer with or without DTT or with the purified reductive pathway components NADPH (1 mM), human Trx1 (25 μM), human TrxR1 (16.25 nM), G6P (1.25 mM), G6PDH (10 U/ml). The SP cells were pelleted by centrifugation at 14,800 × g for 1 min and then incubated with 500 μl PBS containing 25 mM NEM to block free thiols. The samples were washed twice using PBS (1 ml) to remove excess NEM. The cells were lysed on ice with lysis buffer (Tris-HCl (50 mM) containing NaCl (150 mM), EDTA (2 mM), PMSF (0.5 mM), and Triton X-100 (1% v/v)) for 10 min. The lysate was centrifuged at 14,800 × g at 4°C for 10 min. The supernatant was removed and boiled with SDS (2% (w/v)) (VWR chemicals) to denature the protein. Denatured samples were incubated at RT with TCEP (10 mM) to break existing disulfide bonds followed by a 1.5 h incubation with 4-acetamido-4’-maleimidylstilbene-2, 2’-disulfonic acid (20 mM) (AMS) (Invitrogen) in the dark. The samples were separated by SDS-PAGE and a western blot performed with antibody to V5-tag to detect different redox state of ERp57.

### Western blotting

Following SDS-PAGE, proteins were transferred to nitrocellulose blotting membrane (GE Healthcare). The membrane was blocked with 3% w/v dried skimmed milk (Marvel) in TBST buffer (Tris-HCl pH 8.0 (150 mM), NaCl (150 mM), Tween-20 (0.5% v/v)) for 1 h. The membrane was incubated in TBST including V5-antibody (Invitrogen, cat#R960-25) for 1 h. After primary antibody incubation, the membrane was incubated in 5 ml TBST with secondary antibody (Fisher Scientific, cat #10751195) in the dark for 45-60 min. The membrane was scanned by Li-Cor Odyssey 9260 Imager.

### Proteinase K treatment

SP cells were incubated on ice with proteinase K (20 μg/ml) (Roche) for 25 min in the presence of CaCl_2_ (10 mM). Proteinase K was inactivated by incubation with PMSF (0.5 mM). Proteinase K and PMSF was removed by washing twice with KHM buffer.

### Preparation of ADAM10 domain construct and in vitro translation

The human ADAM10 construct codes for residues 456-550 and contains the human-β2M signal sequence residues (1-20). The generation of this construct and its transcription and translation were described previously (Robinson, 2019). Essentially, plasmid DNA was used as a template for PCR using an appropriate forward primer that adds a T7 promoter and a reverse primer that lacks a stop codon. The PCR product was transcribed into an RNA template using T7 RNA polymerase. Translations were performed using the Flexi^®^ Rabbit Reticulocyte Lysate System (Promega) supplemented with DTT (10 mM) or ddH20 but in the absence of added G6P. SP-cells were added to a concentration of ∼10^5^ cells per 25 μl translation reaction. Following assembly of components, translation reactions were incubated at 30 °C for 15 min. For post-translational assays, G6P (5 mM) was added following translation or SP-cells were isolated by centrifugation and resuspended in KHM buffer in the presence or absence of a TrxR1 reduction system NADPH (1 mM), human Trx1 (25 μM), human TrxR1 (16.25 nM), G6P (1.25 mM) and G6PDH (1U) or TCEP (1 mM). An additional incubation was carried out in the presence of the TrxR1 reduction system without Trx1. The reactions were stopped after a 60 min incubation by the addition of NEM (final concentration 25 mM) prior immunoisolation, SDS-PAGE and phosphorimage analysis.

### Immunoisolation

SP-cells from the translation reactions were resuspended in 1 ml immunoisolation buffer (IB) (Tris-HCl (50 mM) pH7.5, Triton X-100 (1% v/v), NaCl (150 mM), EDTA (2 mM), PMSF (0.5 mM), Na-azide (0.02 %)) containing 50 μl 10% Protein A Sepharose FF Resin (Generon). After a 30 min incubation the beads were isolated by centrifugation at 16,000 xg for 30s. The supernatant was transferred into a new tube contained 50 μl 10% PAS. Anti-V5 antibody (Invitrogen) was added and the suspension incubated at 4 °C overnight. The beads were pelleted by centrifugation and washed three times wash using 1ml IB buffer. The beads were heated for 5 min at 105 °C with SDS-PAGE loading buffer (Tris/HCl (200 mM) pH 6.8, containing 10% v/v glycerol, 2% (w/v) SDS, 0.1 % (w/v) bromophenol blue). The samples were separated by SDS-PAGE, the gels were fixed in 10% acetic acid and 10% methanol, dried and exposed to a phosphorimager plate. A FLA-7000 bioimager (Fujifilm) was used to scan the plate to obtain the image.

### Sample preparation for mass spectrometry analysis

Samples were prepared for redox proteomics as previously described (van der Reest, Lilla et al., 2018) with some modifications. Microsomes were incubated in the presence of the reductive system as described above either in the presence or absence of Trx1. After 60 min incubation at 30 °C, microsomes were lysed in a buffer containing 4% SDS, 50 mM triethylammonium bicarbonate pH 8.5 and 55 mM unlabelled iodoacetamide (light IAA, ^12^C_2_H_4_INO – Sigma) to alkylate free cysteine thiols. For each experimental replicate, 30 μg of protein were subsequently labelled with stable-isotope labelled iodoacetamide (heavy IAA, ^13^C_2_D_2_H_2_INO, Sigma) to alkylate reversibly oxidised cysteines. Differentially alkylated proteins were precipitated using trichloroacetic acid (TCA) and digested first with Endoproteinase Lys-C (ratio 1:33 enzyme:lysate) for one h, followed by trypsin, overnight (ratio 1:33 enzyme:lysate). The digested peptides from each experiment, and a pool sample, were differentially labelled using TMT10-plex reagent (Thermo Scientific). Fully labelled samples were mixed in equal amount and desalted using a 100 mg Sep Pak C18 reverse phase solid-phase extraction cartridges (Waters).

### LC/MS/MS protocol

TMT-labelled peptides were analysed as previously described (van der Reest et al., 2018) with minor modifications. First peptides were fractionated using high pH reverse phase chromatography on a C18 column (150 × 2.1 mm i.d. - Kinetex EVO (5 μm, 100 Å)) on a HPLC system (Agilent, LC 1260 Infinity II, Agilent). A two-step gradient was applied, from 1–28% B in 42 min, then from 28–46% B in 13 min to obtain a total of 21 fractions for LC/MS/MS analysis.

Fractionated peptides were separated by nanoscale C18 reverse-phase liquid chromatography using an EASY-nLC II 1200 (Thermo Scientific) coupled to an Orbitrap Fusion Lumos mass spectrometer (Thermo Scientific). Elution was carried out using a binary gradient with buffer A (2% acetonitrile) and B (80% acetonitrile), both containing 0.1% formic acid. Samples were loaded with 6 μl of buffer A into a 50 cm fused silica emitter (New Objective) packed in-house with ReproSil-Pur C18-AQ, 1.9 μm resin (Dr Maisch GmbH). Packed emitter was kept at 50 °C by means of a column oven (Sonation) integrated into the nanoelectrospray ion source (Thermo Scientific). Peptides were eluted at a flow rate of 300 nl/min using different gradients optimised for three sets of fractions: 1–7, 8–15, and 16–21. Initial percentage of buffer B (%B) was kept constant for 3 minutes, then a two-step gradient was used, all with 113 min for step one and 37 min for step two. The %B were changed as follows. For F1-7, %B was 2 at the start, 17 at step one, and 26 at step two. For F8-14, %B was 4 at the start, 23 at step one, and 35 at step two. For F15-21, %B was 6 at the start, 27 at step one, and 43 at step two. All gradients were followed by a washing step (95% B) of 10 min followed by a 5 min re-equilibration step at the initial %B of each gradient, for a total gradient time of 168 min. Eluting peptides were electrosprayed into the mass spectrometer using a nanoelectrospray ion source (Thermo Scientific). An Active Background Ion Reduction Device (ESI Source Solutions) was used to decrease air contaminants signal level. The Xcalibur software (Thermo Scientific) was used for data acquisition. A full scan over mass range of 350–1400 m/z was acquired at 60,000 resolution at 200 m/z, with a target value of 500,000 ions for a maximum injection time of 20 ms. Higher energy collisional dissociation fragmentation was performed on the 15 most intense ions, for a maximum injection time of 100 ms, or a target value of 100,000 ions. Peptide fragments were analysed in the Orbitrap at 50,000 resolution.

### MS Data Analysis

The MS Raw data were processed with MaxQuant software (Cox & Mann, 2008) version 1.6.3.3 and searched with Andromeda search engine (Cox, Neuhauser et al., 2011), querying SwissProt (UniProt, 2010). First and main searches were performed with precursor mass tolerances of 20 ppm and 4.5 ppm, respectively, and MS/MS tolerance of 20 ppm. The minimum peptide length was set to six amino acids and specificity for trypsin cleavage was required, allowing up to two missed cleavage sites. MaxQuant was set to quantify on “Reporter ion MS2”, and TMT10plex was chosen as Isobaric label. Interference between TMT channels were corrected by MaxQuant using the correction factors provided by the manufacturer. The “Filter by PIF” option was activated and a “Reporter ion tolerance” of 0.003 Da was used. Modification by light (H(3)NOC(2)) and heavy (HNOCx(2)Hx(2)) iodoacetamide on cysteine residues (carbamidomethylation) were specified as variable, as well as methionine oxidation and N-terminal acetylation modifications, no fixed modifications were specified. The peptide, protein, and site false discovery rate (FDR) was set to 1 %. MaxQuant outputs were analysed with Perseus software version 1.6.2.3 (Cox & Mann, 2008). The MaxQuant output ModSpecPeptide.txt file was used for quantification of cysteine-containing peptides oxidation, whereas ProteinGroup.txt file was used for protein quantification analysis. Peptides with Cys count lower than one were excluded from the analysis, together with Reverse and Potential Contaminant flagged peptides. From the ProteinGroups.txt file, Reverse and Potential Contaminant flagged proteins were removed, as well as protein groups identified with no unique peptides. The TMT corrected intensities of proteins and peptides were normalised to the median of all intensities measured in each replicate. Only cysteine-containing peptides uniquely assigned to one protein group within each replicate experiment, and robustly quantified in three out of three replicate experiments, were normalised to the total protein levels and included in the analysis. To determine significantly regulated cysteine-containing peptides, a Student t-test with a 5% FDR (permutation-based) was applied using normalised reporter ions intensities.

## Acknowledgements

We wish to acknowledge the generosity of all our colleagues for contributing reagents and other members of the Bulleid group for critical reading of the manuscript. This work was funded by the BBSRC (grant ref: BB/P017665 (the Wellcome Trust (grant number 103720), the China Scholarship Council, and Cancer Research UK (A29800 and A17196).

## Author contributions

The work described in this manuscript was conceived and supervised by NJB with contribution from ZC, XC and SZ. The experimental work was carried out by XC, OBVO, MvL, ZC, MAP, PJR, and SL. The data was analysed by NJB, SL, SZ and XC. The manuscript was written by NJB with editing by all authors.

## References

Appenzeller-Herzog C, Riemer J, Christensen B, Sorensen ES, Ellgaard L (2008) A novel disulphide switch mechanism in Ero1alpha balances ER oxidation in human cells. EMBO J 27: 2977–87

Appenzeller-Herzog C, Riemer J, Zito E, Chin KT, Ron D, Spiess M, Ellgaard L (2010) Disulphide production by Ero1alpha-PDI relay is rapid and effectively regulated. Embo J 29: 3318–29

Baker KM, Chakravarthi S, Langton KP, Sheppard AM, Lu H, Bulleid NJ (2008) Low reduction potential of Ero1alpha regulatory disulphides ensures tight control of substrate oxidation. EMBO J 27: 2988–97

Braakman I, Bulleid NJ (2011) Protein folding and modification in the mammalian endoplasmic reticulum. Annu Rev Biochem 80: 71–99

Bulleid NJ, Ellgaard L (2011) Multiple ways to make disulfides. Trends Biochem Sci 36: 485–92

Cao Z, Mitchell L, Hsia O, Scarpa M, Caldwell ST, Alfred AD, Gennaris A, Collet JF, Hartley RC, Bulleid NJ (2018) Methionine sulfoxide reductase B3 requires resolving cysteine residues for full activity and can act as a stereospecific methionine oxidase. Biochem J 475: 827–838

Chakravarthi S, Bulleid NJ (2004) Glutathione is required to regulate the formation of native disulfide bonds within proteins entering the secretory pathway. J Biol Chem 279: 39872–9

Chakravarthi S, Jessop CE, Bulleid NJ (2006) The role of glutathione in disulphide bond formation and endoplasmic-reticulum-generated oxidative stress. EMBO Rep 7: 271–5

Chen W, Helenius J, Braakman I, Helenius A (1995) Cotranslational folding and calnexin binding during glycoprotein synthesis. Proc Natl Acad Sci U S A 92: 6229–33

Cho SH, Porat A, Ye J, Beckwith J (2007) Redox-active cysteines of a membrane electron transporter DsbD show dual compartment accessibility. EMBO J 26: 3509–20

Cox J, Mann M (2008) MaxQuant enables high peptide identification rates, individualized p.p.b.-range mass accuracies and proteome-wide protein quantification. Nat Biotechnol 26: 1367–72

Cox J, Neuhauser N, Michalski A, Scheltema RA, Olsen JV, Mann M (2011) Andromeda: a peptide search engine integrated into the MaxQuant environment. J Proteome Res 10: 1794–805

Cunnea PM, Miranda-Vizuete A, Bertoli G, Simmen T, Damdimopoulos AE, Hermann S, Leinonen S, Huikko MP, Gustafsson JA, Sitia R, Spyrou G (2003) ERdj5, an endoplasmic reticulum (ER)-resident protein containing DnaJ and thioredoxin domains, is expressed in secretory cells or following ER stress. J Biol Chem 278: 1059–66

Dierks T, Dickmanns A, Preusser-Kunze A, Schmidt B, Mariappan M, von Figura K, Ficner R, Rudolph MG (2005) Molecular basis for multiple sulfatase deficiency and mechanism for formylglycine generation of the human formylglycine-generating enzyme. Cell 121: 541–552

Ellgaard L, Sevier CS, Bulleid NJ (2018) How Are Proteins Reduced in the Endoplasmic Reticulum? Trends Biochem Sci 43: 32–43

Hagiwara M, Maegawa K, Suzuki M, Ushioda R, Araki K, Matsumoto Y, Hoseki J, Nagata K, Inaba K (2011) Structural basis of an ERAD pathway mediated by the ER-resident protein disulfide reductase ERdj5. Mol Cell 41: 432–44

Hoseki J, Oishi A, Fujimura T, Sakai Y (2016) Development of a stable ERroGFP variant suitable for monitoring redox dynamics in the ER. Biosci Rep 36

Jansens A, van Duijn E, Braakman I (2002) Coordinated nonvectorial folding in a newly synthesized multidomain protein. Science 298: 2401–3

Jessop CE, Bulleid NJ (2004) Glutathione directly reduces an oxidoreductase in the endoplasmic reticulum of mammalian cells. J Biol Chem 279: 55341–7

Jessop CE, Chakravarthi S, Garbi N, Hammerling GJ, Lovell S, Bulleid NJ (2007) ERp57 is essential for efficient folding of glycoproteins sharing common structural domains. EMBO J 26: 28–40

Kadokura H, Katzen F, Beckwith J (2003) Protein disulfide bond formation in prokaryotes. Annu Rev Biochem 72: 111–35

Lavery GG, Walker EA, Draper N, Jeyasuria P, Marcos J, Shackleton CH, Parker KL, White PC, Stewart PM (2006) Hexose-6-phosphate dehydrogenase knock-out mice lack 11 beta-hydroxysteroid dehydrogenase type 1-mediated glucocorticoid generation. J Biol Chem 281: 6546–51

Lillig CH, Holmgren A (2007) Thioredoxin and related molecules--from biology to health and disease. Antioxid Redox Signal 9: 25–47

Molteni SN, Fassio A, Ciriolo MR, Filomeni G, Pasqualetto E, Fagioli C, Sitia R (2004) Glutathione limits Ero1-dependent oxidation in the endoplasmic reticulum. J Biol Chem 279: 32667–73

Ohishi K, Inoue N, Maeda Y, Takeda J, Riezman H, Kinoshita T (2000) Gaa1p and gpi8p are components of a glycosylphosphatidylinositol (GPI) transamidase that mediates attachment of GPI to proteins. Mol Biol Cell 11: 1523–33

Oka OB, Pringle MA, Schopp IM, Braakman I, Bulleid NJ (2013) ERdj5 is the ER reductase that catalyzes the removal of non-native disulfides and correct folding of the LDL receptor. Mol Cell 50: 793–804

Oliver JD, van der Wal FJ, Bulleid NJ, High S (1997) Interaction of the thiol-dependent reductase ERp57 with nascent glycoproteins. Science 275: 86–8

Palde PB, Carroll KS (2015) A universal entropy-driven mechanism for thioredoxin-target recognition. Proc Natl Acad Sci U S A 112: 7960–5

Poet GJ, Oka OB, van Lith M, Cao Z, Robinson PJ, Pringle MA, Arner ES, Bulleid NJ (2017) Cytosolic thioredoxin reductase 1 is required for correct disulfide formation in the ER. EMBO J 36: 693–702

Ponsero AJ, Igbaria A, Darch MA, Miled S, Outten CE, Winther JR, Palais G, D’Autreaux B, Delaunay-Moisan A, Toledano MB (2017) Endoplasmic Reticulum Transport of Glutathione by Sec61 Is Regulated by Ero1 and Bip. Mol Cell 67: 962–973 e5

Rishavy MA, Usubalieva A, Hallgren KW, Berkner KL (2011) Novel insight into the mechanism of the vitamin K oxidoreductase (VKOR): electron relay through Cys43 and Cys51 reduces VKOR to allow vitamin K reduction and facilitation of vitamin K-dependent protein carboxylation. J Biol Chem 286: 7267–78

Roberts BS, Babilonia-Rosa MA, Broadwell LJ, Wu MJ, Neher SB (2018) Lipase maturation factor 1 affects redox homeostasis in the endoplasmic reticulum. EMBO J 37

Robinson P, Kanemura, S, Cao, X, Bulleid, NJ (2019) Protein secondary structure determines the temporal relationship between folding and disulfide formation. BioRxiv 564740

Schwertassek U, Balmer Y, Gutscher M, Weingarten L, Preuss M, Engelhard J, Winkler M, Dick TP (2007) Selective redox regulation of cytokine receptor signaling by extracellular thioredoxin-1. EMBO J 26: 3086–97

Sevier CS, Qu H, Heldman N, Gross E, Fass D, Kaiser CA (2007) Modulation of cellular disulfide-bond formation and the ER redox environment by feedback regulation of Ero1. Cell 129: 333–44

Shepherd C, Oka OB, Bulleid NJ (2014) Inactivation of mammalian Ero1alpha is catalysed by specific protein disulfide-isomerases. Biochem J 461: 107–13

Tavender TJ, Springate JJ, Bulleid NJ (2010) Recycling of peroxiredoxin IV provides a novel pathway for disulphide formation in the endoplasmic reticulum. EMBO J 29: 4185–97

Tie JK, Jin DY, Stafford DW (2014) Conserved loop cysteines of vitamin K epoxide reductase complex subunit 1-like 1 (VKORC1L1) are involved in its active site regeneration. J Biol Chem 289: 9396–407

Tie JK, Jin DY, Tie K, Stafford DW (2013) Evaluation of warfarin resistance using transcription activator-like effector nucleases-mediated vitamin K epoxide reductase knockout HEK293 cells. J Thromb Haemost 11: 1556–64

Tsunoda S, Avezov E, Zyryanova A, Konno T, Mendes-Silva L, Pinho Melo E, Harding HP, Ron D (2014) Intact protein folding in the glutathione-depleted endoplasmic reticulum implicates alternative protein thiol reductants. Elife 3: e03421

UniProt C (2010) The Universal Protein Resource (UniProt) in 2010. Nucleic Acids Res 38: D142–8

Ushioda R, Hoseki J, Araki K, Jansen G, Thomas DY, Nagata K (2008) ERdj5 is required as a disulfide reductase for degradation of misfolded proteins in the ER. Science 321: 569–72

van der Reest J, Lilla S, Zheng L, Zanivan S, Gottlieb E (2018) Proteome-wide analysis of cysteine oxidation reveals metabolic sensitivity to redox stress. Nat Commun 9: 1581

van Lith M, Tiwari S, Pediani J, Milligan G, Bulleid NJ (2011) Real-time monitoring of redox changes in the mammalian endoplasmic reticulum. J Cell Sci 124: 2349–56

Wang M, Kaufman RJ (2016) Protein misfolding in the endoplasmic reticulum as a conduit to human disease. Nature 529: 326–35

Wilson R, Allen AJ, Oliver J, Brookman JL, High S, Bulleid NJ (1995) The translocation, folding, assembly and redox-dependent degradation of secretory and membrane proteins in semi-permeabilized mammalian cells. Biochemical Journal 307 (Pt 3): 679–87

